# The Effectiveness of China’s National Forest Protection Program and National-level Nature Reserves, 2000 to 2010: PREPRINT

**DOI:** 10.1101/000893

**Authors:** Guopeng Ren, Stephen S. Young, Lin Wang, Wei Wang, Yongcheng Long, Ruidong Wu, Junsheng Li, Jianguo Zhu, Douglas W. Yu

## Abstract

There is profound interest in knowing the degree to which China’s institutions are capable of protecting its natural forests and biodiversity in the face of economic and political change. China’s two most important forest protection policies are its National Forest Protection Program (NFPP) and its National-level Nature Reserves (NNRs). The NFPP was implemented in 17 provinces starting in the year 2000 in response to deforestation-caused flooding. We used MODIS data (MOD13Q1) to estimate forest cover and forest loss across mainland China, and we report that 1.765 million km^2^ or 18.7% of mainland China was covered in forest (12.3%, canopy cover > 70%) and woodland (6.4%, 40% ≤ canopy cover < 70%) in 2000. By 2010, a total of 480,203 km^2^ of forest + woodland was lost, amounting to an annual deforestation rate of 2.7%. The forest-only loss was 127,473 km^2^, or 1.05% annually. The three most rapidly deforested provinces were outside NFPP jurisdiction, in the southeast. Within the NFPP provinces, the annual forest + woodland loss rate was 2.26%, and the forest-only rate was 0.62%. Because these loss rates are likely overestimates, China appears to have achieved, and even exceeded, its NFPP target of reducing deforestation to 1.1% annually in the target provinces. We also assemble the first-ever polygon dataset for China’s forested NNRs (n = 237), which covered 74,030 km^2^ in 2000. Conventional unmatched and covariate-matching analyses both find that about two-thirds of China’s NNRs exhibit effectiveness in protecting forest cover and that within-NNR deforestation rates are higher in provinces that have higher overall deforestation.

## Introduction

China covers one of the greatest ranges of ecological diversity in the world, in total containing perhaps 10% of all species living on Earth (1). Maintaining much of this biodiversity is a substantial cover of natural forest, but China’s biodiversity, ecosystem services, and natural landscapes are being degraded at a rapid rate, due to a combination of swift economic growth and institutional constraints (2–5). With the decline of its natural forests, China has witnessed a deterioration in biodiversity, with at least 200 plant species lost, 15-20% of China’s higher plant species endangered, 233 vertebrate species on the edge of extinction, and more than 61% of wildlife species suffering habitat loss (1, 6, 7). On the other hand, China has made important political commitments to conservation as a signatory of the major international biodiversity treaties such as the Convention on Biological Diversity (CBD) and the Convention on International Trade in Endangered Species of Wild Fauna and Flora (CITES), as well as implementing many domestic conservation laws and regulations, establishing a Ministry of Environmental Protection, and targeting conservation programs for charismatic species such as panda, tiger, multiple primates, and cranes.

Arguably the most important class of conservation measures in China has been its forest-protection policies. During the 1970s to 1990s, with weak forest-protection laws and enforcement, the annual logging rate is estimated to have ranged from 2.67% to 3.36%, and the annual deforestation rate from 1994 to 1996 is estimated to have been as high as 3.0% (8). This extensive deforestation contributed to severe water and wind erosion of soil, so that by the early 1990s, approximately 38% of China’s total land area was considered badly eroded (9–11). In particular, deforestation and cultivation on steep slopes in Yunnan and western Sichuan provinces led to the devastating 1998 flood in the Yangtze River basin (12–15).

In response, mainland China piloted the Natural Forest Protection Program (NFPP) in 1998 across 12 of 31 total provinces to prevent the further loss of forest cover and to expand afforestation. In 2000, the NFPP (2000-2010) was formally implemented across 17 provinces in three regions, (1) the upper reaches of the Yangtze River (Tibet, Yunnan, Sichuan, Guizhou, Chongqing, and Hubei provinces), (2) the upper and middle reaches of the Yellow River (Qinghai, Gansu, Ningxia, Shaanxi, Shanxi, Henan, and Inner Mongolia), and (3) the key state-owned forestry regions (Heilongjiang, Inner Mongolia, Jilin, Xinjiang, and Hainan) (Fig. 1, Fig. 2 inset). A total of 96.2 billion RMB (∼ US$14.5 billion) was budgeted to implement the NFPP from 2000 through 2010 (9, 10, 16–18), and the final expenditure (2000-2010) ended up growing to 118.6 billion RMB (19, 20).

**Fig. 1.**
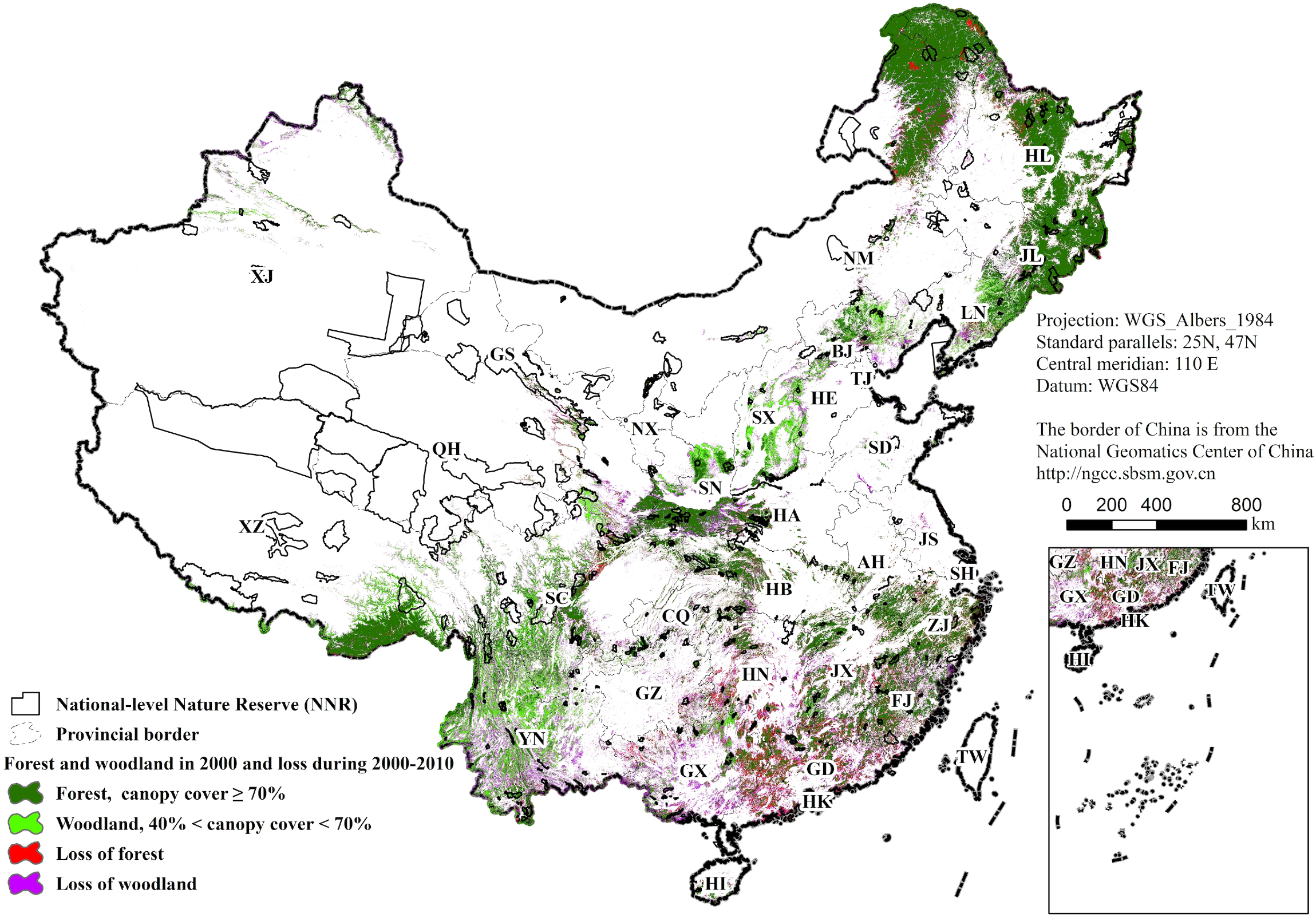
Map of forest and woodland coverage over mainland China in year 2000 and loss during 2000-2010. AH-Anhui, BJ-Beijing, CQ-Chongqing, FJ-Fujian, GD-Guangdong, GS-Gansu, GX-Guangxi, GZ-Guizhou, HA-Henan, HB-Hubei, HE-Hebei, HI-Hainan, HL-Heilongjiang, HN-Hunan, JL-Jilin, JS-Jiangsu, JX-Jiangxi, LN-Liaoning, NM-Inner Mongolia, NX-Ningxia, QH-Qinghai, SC-Sichuan, SD-Shandong, SH-Shanghai, SN-Shaanxi, SX-Shanxi, TJ-Tianjin, XJ-Xinjiang, XZ-Tibet, YN-Yunnan, ZJ-Zhejiang. A high-resolution version is in SI Text 3.

**Fig. 2.**
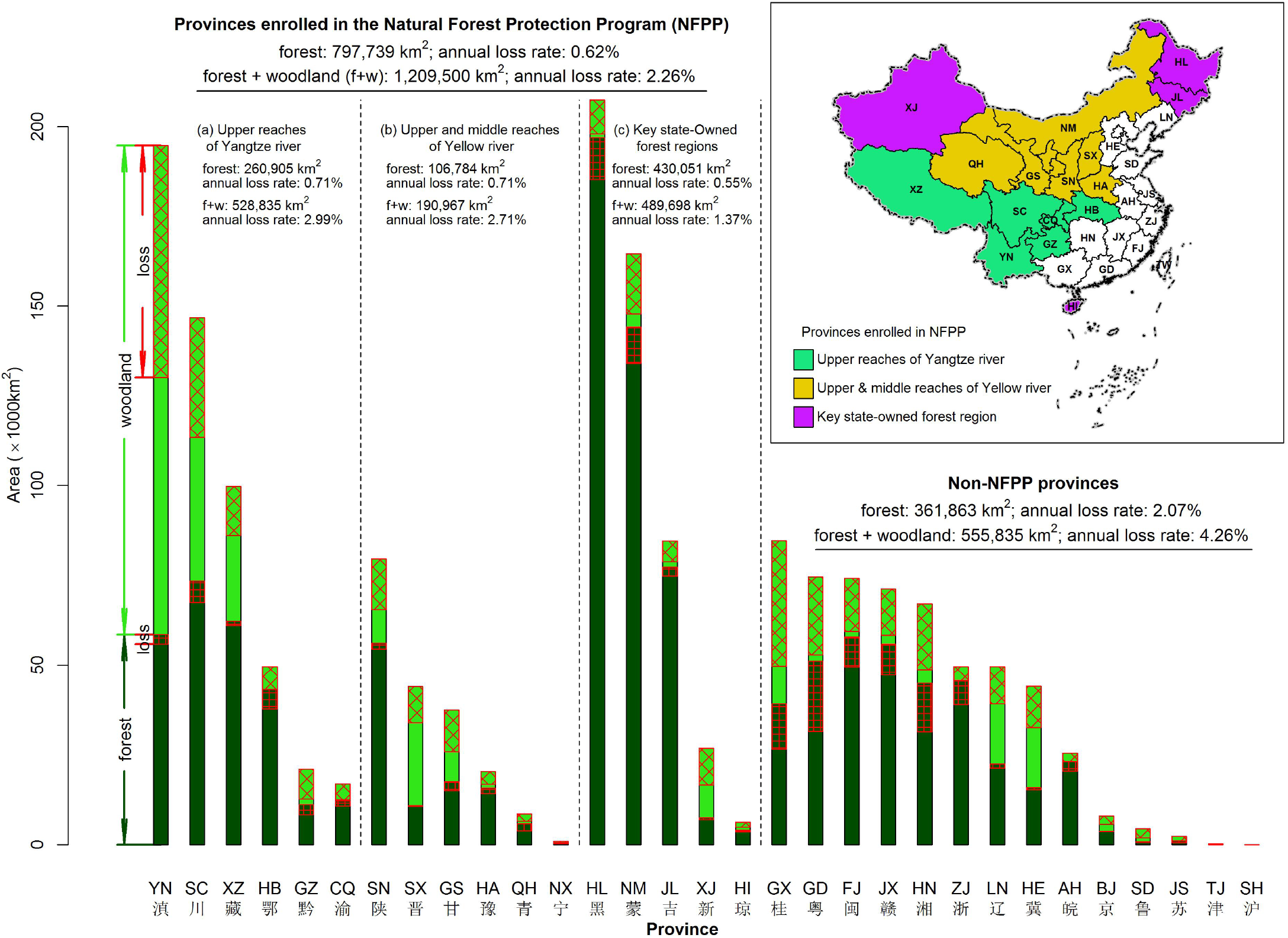
Forest and woodland coverage over mainland China in year 2000 and loss during 2000-2010, subdivided by province and by National Forest Protection Program (NFPP) status. Inset: NFPP-enrolled provinces.

The NFPP’s main objective related to reducing deforestation was to reduce commercial timber extraction by 62.1% (i.e. from 13.518 million m^3^ in 1997 to 1.128 million m^3^ by the year 2000 in the upper reaches of the Yangtze River and the upper and middle reaches of the Yellow River; and from 18.532 million m^3^ in 1997 to 11.017 million m^3^ by the year 2003 in the key state-owned forestry region). This reduction included a ban on commercial logging on 0.445 million km^2^ (17, 18). Given that the annual logging rate during 1994-1996 across mainland China was 3.0% (8), to achieve a 62.1% reduction in logging in the NFPP zone implies that the annual deforestation rate within the NFPP zone needed to be reduced to 1.1% (=3.0% X (1-62.1%)) during 2000-2010. To judge the effectiveness of the NFPP, we can compare this target against the achieved deforestation rate.

China also protects natural forest within its nature reserve (NR) network. Starting with its first NR in 1956, China has established 2,669 NRs as of year 2012, covering 14.9% of China’s land area (1.43 million km^2^) (21). 363 of them are National-level Nature Reserves (NNRs), which are meant to cover the most important ecosystems and areas for biodiversity in China and consequently receive the highest level of protection and state budget. NNRs account for 62.9% of the area of all NRs (21). This level of coverage puts China close to the upper range of the global mean coverage (10.1-15.5% of a nation’s land area) (22, 23), although a large fraction is accounted for by a handful of reserves on the plateaus of Tibet, Xinjiang, and Qinghai (23) (Fig. 1).

However, the effectiveness of China’s NFPP and NNR programs in protecting natural forest is disputed (24–26). Even the precise borders of the NNRs have been mostly unknown to the public, and estimates of China’s forest cover between 2000 and 2010 have varied by a factor of 1.7 (1,209,000-2,054,056 km^2^), in part due to varying definitions (27–30). Moreover, because economic development typically takes precedence over biological conservation at local governance levels (6), it is widely reported that natural forest continues to be cleared or converted to plantations, especially rubber and fruit trees (5, 31–34), which directly contravenes the NFPP’s primary policy goal of reducing the loss of standing forest.

Given China’s high absolute endowment of biodiversity and forest cover and the profound domestic and international interest in knowing the degree to which modern China’s governing institutions are capable of protecting biodiversity, (1) we mapped annual forest cover over the whole of mainland China from 2000 through 2010, using a uniform remote sensing dataset (231.7 m resolution MODIS product, MOD13Q1) and the *randomForest* algorithm (35–38), (2) we quantified deforestation across mainland China with a three-year moving window in the NFPP-era from 2000 to 2010, and (3) we used both conventional unmatched and covariate-matching analyses (39, 40) to estimate how much closed forest in the NNRs avoided deforestation during the logging-ban era.

Estimates of forest-cover loss provide rare quantitative measures of the effectiveness of China’s nature-protection institutions, due to the NFPP’s public, quantitative deforestation reduction targets, and because NNR status prohibits deforestation outright.

## Results

### Forest cover and loss 2000-2010

In 2000, 1.765 million km^2^ or 18.7% of mainland China was covered in forest (12.3%, defined here as canopy cover ≥ 70%) or woodland (6.4%, 40% ≤ canopy cover < 70%) (Fig. 1). Consistent with previous reports (41–43), most of the forest and woodland cover was located in the northeast, followed by the southwest and then the southeast; just five provinces accounted for 46% of forest and woodland cover (Heilongjiang *HL*, Yunnan *YN*, Inner Mongolia *NM*, Sichuan *SC*, and Tibet *XZ*) (Fig. 2).

From 2000 to 2010, we recorded forest loss and ignored any subsequent afforestation because our goal is to measure the effectiveness of the NFPP and NNRs in reducing and preventing any loss, respectively, of already standing forest. Also, in China, most afforestation and reforestation is to monoculture plantations, which provide less biodiversity, carbon storage, and other ecosystem services (31, 44–46).

We define *narrow-sense deforestation* (δ_forest_) as clearance of forest to non-wooded land, and *broad-sense deforestation* (δ_forest+woodland_) is defined as clearance of forest + woodland to non-wooded land. δ_forest+woodland_ is a less reliable measure, due to a higher probability of mis-registration (**Methods, SI Text 1 for further details**). For the same reason, we did not attempt to record whether forest converted to woodland or woodland to forest.

Most narrow-sense deforestation occurred in the southeast, with the top three provinces of Guangdong *GD*, Hunan *HN*, and Guangxi *GX* (all non-NFPP) alone accounting for 36.0% of δ_forest_ (Figs. 1, 2). The next two provinces with the highest δ_forest_ values, Heilongjiang *HL* and Inner Mongolia *NM*, are designated key state-owned forest regions within the NFPP, have the highest proportion of forest cover in the country and accounted for 17.1% of δ_forest_ from 2000 to 2010 (Fig. 2). For the 27 mainland provinces with forest area larger than 1000 km^2^ in 2000, provincial forest cover did not explain deforestation rate (linear regression, R^2^ = 0.06, F_1,25_ = 1.63, p = 0.23).

From 2000 to 2010, δ_forest+woodland_ was 480,203 km^2^, or 27.2% of the original forest + woodland cover, resulting in 1.285 million km^2^ or 13.6% of mainland China remaining under original forest or woodland in 2010. The top five provinces for deforestation were all in the south and southwest, again including the same three non-NFPP provinces, Guangdong *GD*, Hunan *HN*, and Guangxi *GX*, plus two NFPP provinces, Yunnan *YN* and Sichuan *SC*, which collectively were responsible for 47.3% of δ_forest+woodland_ (Figs. 1, 2).

### Testing NNR effectiveness

We evaluated the effectiveness of the NNRs at the individual reserve level and collectively at the provincial level, as NNRs are managed at both scales. At each level we used both *unmatched* and *matched* (=‘covariate matching’) sampling methods (39, 40) to estimate δ_forest_ by sampling pixels that were forest in the year 2000 and determining if these pixels were deforested (neither forest nor woodland) in 2010. In brief, the unmatched method randomly samples a set number of forested pixels in year 2000 inside the NNR and again outside the NNR (10-100 km from the NNR border) and compares how many pixels inside and outside remained forested in 2010. The matched sampling method also compares a set number of pixels inside and outside of the NNRs, but pairs of inside and outside pixels are matched by characteristics such as elevation, slope, and distance to forest edge. The matched method is more effective at isolating the effect of reserve status *per se* because it controls for nature reserves being more likely to have been sited in more remote and less productive areas, thus deriving part of their protection from greater inaccessibility.

237 of China’s 363 NNRs contained sufficient natural forest both inside and outside their borders to be evaluated (**Methods**). The 237 NNRs, which enclosed a total area of 74,030 km^2^ of forest in the year 2000, prevented 4,073 km^2^ of forest loss (Fig. 3a), as evaluated by the unmatched approach, or prevented 3,148 km^2^ (Fig. 3b), as evaluated by the matched approach. The two methods largely concurred; the unmatched method judged 167 NNRs to be effective (i.e. prevented a statistically significant amount of forest loss, p < 0.05, paired t-test with *fdr* correction, Supplementary Table S1), and the matched method found the same for 158 NNRs. 137 NNRs were found effective under both methods, and 188 NNRs were judged effective by at least one method (**SI Text 2** for per-NNR results).

**Fig. 3.**
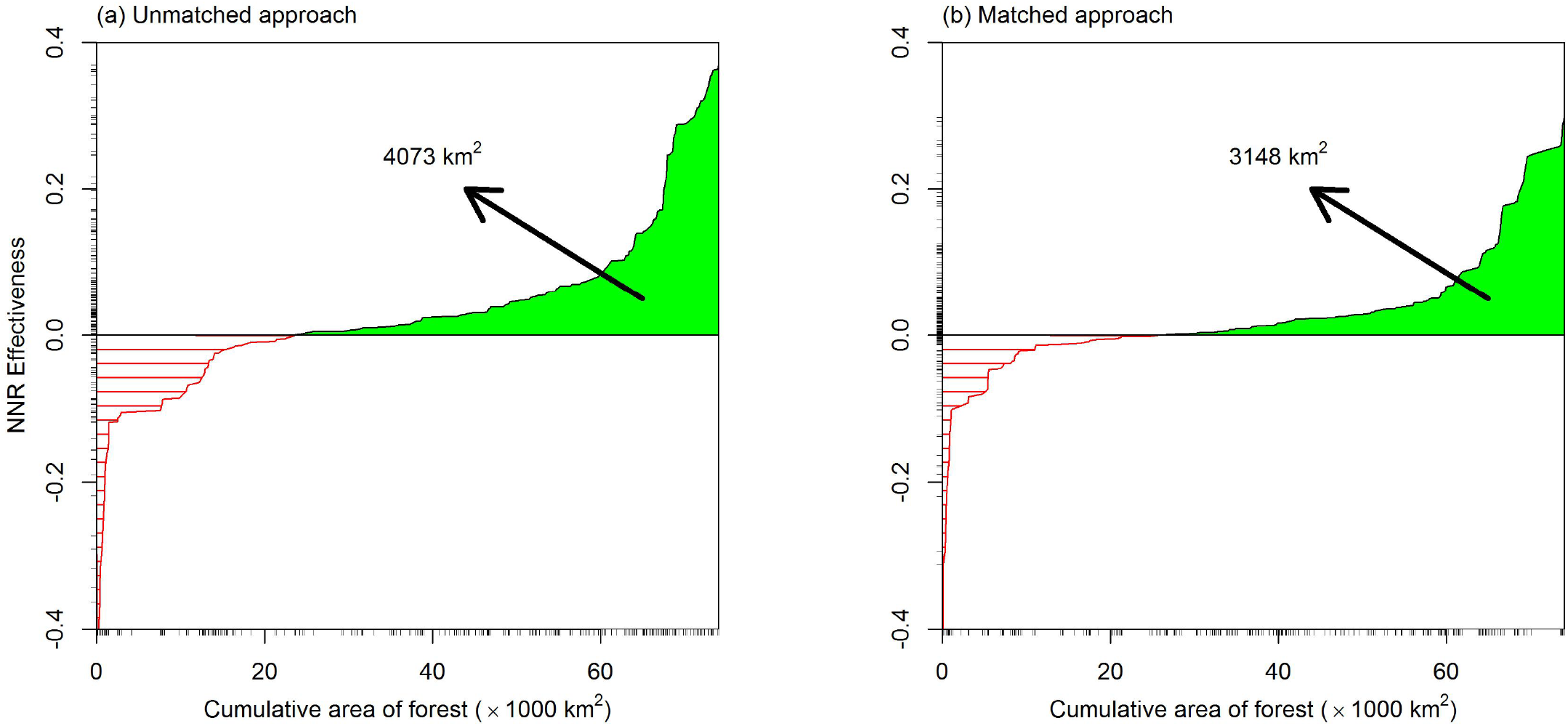
Effectiveness of National-level Nature Reserves (NNRs) in protecting forest cover during 2000-2010. Calculation details are described in **Methods: NNR Effectiveness**. NNRs are ordered from lowest (negative) to highest (positive) effectiveness. Rug lines (small ticks) along the X-axis indicate each NNR’s contribution to total forest cover. Rug lines along the Y-axis indicate each NNR’s effectiveness. (A) Unmatched method. (B) Matched method.

We also measured the pooled effectiveness of NNRs at the provincial governance level, including all provinces with forest cover greater than 1000 km^2^, thus excluding Shandong, Jiangsu, Tianjin and Shanghai for lack of 1000 km^2^ forest cover, and Ningxia where forest cover was almost exclusively located in NNRs, preventing comparison. Pooled NNRs in 23 of 26 provinces were effective (p < 0.05, paired t-test with *fdr* correction) in protecting any forest cover, as evaluated by either approach (unmatched: 12 of 26 provinces; matched: 17 of 26) (Fig. 4). As a natural result of comparing inside-NNR pixels against those outside, the higher the deforestation rate in the province, the more effective did the NNRs appear to be in that province (linear regression, unmatched: R^2^ = 0.49, F_1,24_ = 23.44, p < 0.001; matched: R^2^ = 0.71, F_1,24_ = 59.81, p < 0.001) (Fig. 4).

**Fig. 4.**
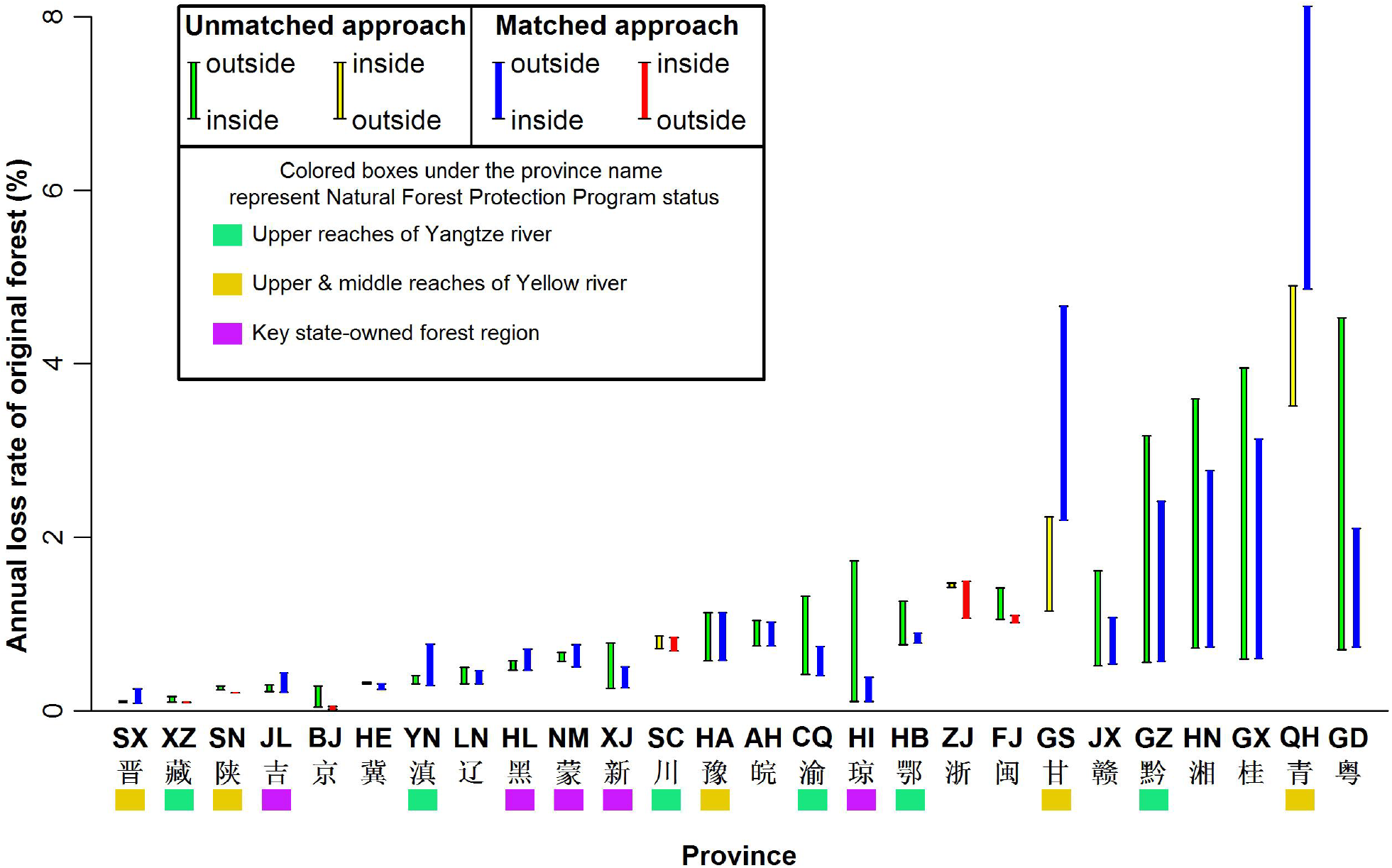
Effectiveness of National-level Nature Reserves (NNRs) in protecting forest cover during 2000-2010, pooled by province. Provinces are aligned from left to right by each province’s deforestation rate. For each province, effectiveness is estimated using matched and unmatched methods. For blue and green bars, the deforestation rate inside the NNRs is lower than outside the NNRs. For red and yellow bars, the deforestation rate inside the NNRs is higher than outside the NNRs. Most provinces show a lower deforestation rate inside the NNRs, indicating a protective effect of the province’s NNRs. Color bars below each province indicate National Forest Protection Program (NFPP) status (color scheme follows Figure 2 inset).

## Discussion

Although the loss of biodiversity and natural forests are global concerns, the institutions for managing and conserving these resources mostly originate domestically, especially in large countries like China, which have endogenously developed legal and political systems. During 2000-2010, we estimate that already extant, mostly natural forest cover declined from 12.3% to 10.9% of mainland China, and original forest + woodland cover declined from 18.7% to 13.6%. From 2000 to 2010, this translates to a 1.05% annual loss rate and a half-life of 65 years for forest only, and to a 2.84% annual loss rate and a 24-year half-life for forest + woodland.

Most of the narrow-sense deforestation (δ_forest_) occurred in the 14 non-NFPP provinces (annual rate 2.07% during 2000-2010), especially in southeastern China (Figs. 1 and 2), while the annual rate across the 17 NFPP provinces was 3.3 times lower, at 0.62%. Moreover, within the NFPP provinces, most of the narrow-sense deforestation was concentrated in two provinces, Heilongjiang *HL* and Inner Mongolia *NM*, in northeastern China (Fig. 2), which were designated in the NFPP as “key state-owned forest regions”. Logging was not as strictly banned in this region as in the watersheds of the Yangtze and Yellow rivers.

Broad-sense deforestation (δ_forest+woodland_) was similarly higher in the non-NFPP provinces, especially in southeastern China (annual rate 4.26% during 2000-2010), than in the NFPP provinces (2.26%). However, two NFPP provinces, Yunnan *YN* and Sichuan *SC*, through which the Yangtze river passes and which house a high fraction of China’s biodiversity (23), also show high levels of δ_forest+woodland_ (Fig. 2).

Although not a focus of our analysis, several studies suggest that an important driver of forest loss across all provinces is replacement by tree plantations (31, 32, 34, 42, 47), especially *Eucalyptus*, to meet high demand for wood products (48–50). In Yunnan *YN,* Guangdong *GD* and Hainan *HI*, which have tropical to subtropical climates, rubber plantations are also an important driver (34, 51). Northwest Yunnan *YN* and western Sichuan’s *SC* woodland loss is more difficult to explain, although there was pressure on forests from increases in tourism and living standards, which was met in part by the construction of traditional, wooden guesthouses and the use of wood for fuel in the Tibetan regions of Yunnan, Sichuan, and Qinghai (52). It also appears that some 1970s-era planted forests in Sichuan and Yunnan were given permission to be logged.

Of course, provinces differ in many more ways than whether they were included in the NFPP, so we cannot unambiguously attribute lower deforestation in the NFPP provinces to the success of that program. However, we can observe that our estimates for the 2000-2010 annual rates of broad- and narrow-sense deforestation in the NFPP provinces (δ_forest+woodland_ = 2.26%, δ_forest_ = 0.62%) approach and more than achieve, respectively, the target of reducing NFPP-zone annual deforestation to 1.1% (Fig. 2). In addition, comparison with Hansen et al.’s (43) recent Landsat-based analysis of global forest cover gives us further reason to believe that China more than achieved their NFPP reduction.

Hansen et al. (43) used 30-meter-resolution Landsat imagery to analyze forest change between 2000 and 2012 at the global scale. Their estimates for China are consistent with ours in terms of year-2000 forest cover (this study: 1.765 million km^2^ 40-100% tree cover; Hansen et al.: 1.702 million km^2^ 26-100% tree cover) and the spatial distribution of cover and subsequent loss (Figs. 1 from the two studies). However, Hansen et al. found much lower deforestation rates than we did, which is likely explained by the smaller pixel sizes afforded by Landsat images (30 × 30 m = 900 m^2^), relative to MODIS (231.7 × 231.7 m = 53685 m^2^). Partial deforestation in a MODIS pixel is sometimes scored as whole-pixel deforestation, leading to overestimates. Most of the deforestation in China captured by Hansen et al. (43) was in small patches, often a few Landsat pixels in size. Where there are large contiguous patches of deforestation such as in Heilongjiang, our estimates are much closer to theirs. A second possible explanation is that we count forest loss each year, and if forest is cut in 2001 and replanted to fast growing rubber or eucalyptus, within a short time, the newly reforested area will have a signature of forest cover and could be missed by their annual percent tree cover filter. Quantifying the contributions of these two factors will need analysis of the EarthEngine Landsat data, when they become available.

The key takeaway is that Hansen et al.’s analysis strongly suggests that our analysis is conservative and that China therefore likely overachieved its NFPP deforestation reduction target for both forest and woodland ecotypes.

Our analysis of China’s forested NNRs lets us conclude that NNR designation also successfully *reduced* deforestation in most provinces and in approximately two-thirds of NNRs (Figs. 3, 4) but have not collectively *prevented* deforestation outright. Moreover, total avoided deforestation (estimated range 3148-4073 km^2^, Fig. 3) amounts to around just half the area of Shanghai municipality (6340 km^2^). This small number derives from the simple fact that China’s NNR areal coverage is biased away from forest ecosystems (Fig. 1) (23, 53).

On the other hand, our analysis of NNR effectiveness is conservative for two reasons. We did not differentiate natural (e.g. fire, insects) from anthropogenic causes of forest loss. In particular, Sichuan’s *SC* NNRs were found to be ineffective (Fig. 4), but this is partly due to landslides from the 2008 Wenchuan earthquake, which caused forest loss within some NNRs (54). Also, we only analyzed national-level reserves and had to ignore provincial, municipal and county-level reserves due to missing polygon data. Thus, some of our outside-NNR pixels might have been in these lower-level reserves, which, if protected, would have reduced the inferred effectiveness of the NNRs. We also emphasize that we did not analyze woodland loss alone, due to limitations of our data, and it is possible that NNR effectiveness could be lower in this forest-cover class.

The effectiveness of the NNRs varies across provinces (Figure 4). In part, this is the natural consequence of using matching methods; an NNR that successfully protects forest cover inside its borders will appear more effective the greater the deforestation rate outside its borders. Thus, estimated effectiveness is greater in non-NFPP provinces, particularly in the deforestation hotspot of Guangdong *GD*, Hunan *HN*, and Guangxi *GX* (**Results**, Fig. 4). Importantly, we observe that the deforestation rate *inside* NNRs is positively correlated with deforestation in the province as a whole (linear regression, unmatched: R^2^ = 0.49, F_1,24_ = 23.44, p < 0.001; matched: R^2^ = 0.71, F_1,24_ = 59.81, p < 0.001). We speculate that causality runs in the direction of generally laxer forest-protection governance at the provincial level (e.g. not being an NFPP province) leading to lower levels of protection inside NNRs as well.

Hainan *HI* provides a clear example of how the matched method can correct for the tendency of the unmatched method to overestimate NNR effectiveness (Fig. 4) (40). Hainan’s forested NNRs are all located at higher elevations, which are unsuitable for rubber and other tropical plantations, and are directly surrounded by converted lowlands (26). An interesting counterexample is given by NNRs in Qinghai *QH* and Gansu *GS*, which were deemed *more* effective by the matched method than by the unmatched method (Fig. 4). This is because Qinghai’s and Gansu’s NNRs happen to be located on flatter and lower altitude sites, rendering them more vulnerable than neighboring forests. When the matched method is used to compare with unprotected sites of equal vulnerability, these NNRs are deemed to be effective. We note that, overall, deforestation rates in these two grassland-dominated provinces are some of the highest in China, despite being NFPP provinces.

In summary, our study provides the first assessment of China’s two most important forest-protection policies. Since the introduction of the NFPP in 2000, at what could be considered the beginning of the modern era of environmental protection in China, the annual forest loss rate has declined to, at most, a moderately low 1.05% across the country, but the annual forest + woodland loss rate is higher in non-NFPP and in two NFPP provinces, Sichuan *SC* and Yunnan *YN*, which are biodiversity hotspots in China. Our results are consistent with many geographically focused analyses in China (31, 32, 34, 42, 47, 51, 55) reporting that natural forests are being replaced by plantations, with the added nuance that it appears to be mostly woodland that is being replaced. The NFPP has been renewed for 2011-2020, and it will be instructive to continue monitoring the performance of this second stage of the NFPP (19, 20).

The reason that NNRs in China have not been comprehensively assessed earlier is because most nature reserves in China, even the national-level reserves, do not have published borders. Establishing a public and standardized database of all nature-reserve borders is a key conservation priority for China. Another urgent priority is to establish or upgrade nature reserves in the south and east of China, where coverage is clearly inadequate and where deforestation rates are highest (23). Finally, it is instructive to compare China with Brazil, which in 2007 implemented a series of institutional and legal reforms, including a near-real-time satellite monitoring system based on MODIS data, to detect and deter deforestation in the Brazilian Amazon. It is estimated that from 2007 to 2011, these reforms reduced expected deforestation by 59-75%, with no reduction in agricultural production (56). Like Brazil, China’s demonstrated ability to implement complex and large-scale policy mechanisms, combined with an advanced technological and scientific infrastructure, provides a clear opportunity for China to continue improving its protection of environmental quality, wildlife, and forests across its entire landscape.

## Materials and Methods

### Remote sensing data

The MOD13Q1 dataset has 12 layers, including a 16-day composite of red (620–670 nm), near infrared (NIR: 841–876 nm), and mid-infrared (MIR: 2105–2155 nm) reflectance, Normalized Difference Vegetation Index (NDVI), and Enhanced Vegetation Index (EVI) at 231.7 m resolution, and it is designed to provide consistent spatial and temporal comparison of vegetation conditions (57). We employed MOD13Q1 to map land cover and deforestation over mainland China from 2000-2010. Low-quality pixels due to clouds and shadows were filled by the mean values of the previous and the next scenes.

### Classification scheme and training data definition

Using Arc2Earth software (http://www.arc2earth.com/), we were able to synchronize our MODIS data with high-resolution images in Google Earth (http://www.google.com/earth), and then in ArcGIS (Ver. 10.0, ESRI) we digitized and visually assigned 13869 polygons (minimal edge > 500 m) to three classes: (1) forest (tree canopy cover ≥ 70%), (2) woodland (tree canopy cover < 70% and tree + shrub cover ≥ 40%), and (3) non-wooded land (tree cover < 40%, including open shrub, grassland, farmland, urban, open water, etc.).

### Land-cover mapping

Because of the large quantity of MODIS data, we partitioned mainland China into 34 contiguous regions to minimize the number of different MODIS tiles needing to be stitched together. Using 23 scenes of MOD13Q1 product per year and the Google Earth training data, we used the *randomForest* algorithm to classify annual land cover over mainland China from 2000 through 2010 (36, 37, 58). For each year, producer’s accuracies in all 34 regions were ≥ 92%.

### Deforestation detection

We used three-year moving windows to adjudge forest loss (forest to non-forest, non-woodland). Only if the first year of a window was forest and the next two years were non-forest and non-woodland was a pixel classed as having been deforested in that window, but if any two of the three years were classed as forest, then the pixel remained closed forest in this window. Our three-year moving window is useful for cases where forest cover is distributed over small patches relative to MODIS pixel size (231.7 m), as we found for one NNR in Gansu. Mis-registration between consecutive years might make some patches appear to get deforested and then later reforested, and our methodology would classify this pixel as deforested because we do not consider reforestation. The three-year window minimizes this mis-registration effect.

To assess change-detection accuracy, we sampled 200 random deforestation pixels in four regions (two in southern China, one in eastern China and one in northeastern China), which could be independently classed as deforested or not using high-resolution imagery in Google Earth. User accuracies were ≥ 90% in all four regions.

### Terrain data

The Shuttle Radar Topography Mission (SRTM) dataset was downloaded from the USGS (http://srtm.usgs.gov/index.php) and re-projected to the Albers conic equal area projection for all 34 regions at 90 m resolution. Elevation, slope and a topographic position index (TPI) were calculated from the 90 m resolution SRTM. The TPI of a focal pixel was defined as the difference between its elevation and the mean elevation of all pixels in an 11 × 11 grid centered on the focal pixel. TPI identifies pixels that are either higher (peaks) or lower (gorges) than the surroundings and thus captures local inaccessibility.

### NNR data

The Chinese Research Academy of Environmental Sciences delineated and merged boundary polygons of all NNRs on behalf of the Ministry of Environmental Protection (MEP). Data sources are the NNR Master Plans finished and reported to the MEP by each NNR under the Regulations on Nature Reserves of the People’s Republic of China. These polygons are currently the best available data on NNR borders, but boundary accuracies do vary because about 20% of the NNRs still have low technological capacity for mapping.

### NNR effectiveness

At the end of 2012, there were 363 NNRs in mainland China. 241 of them contained at least 10 km^2^ of forest cover and were distributed over 29 provinces. Of these, 4 NNRs in two provinces (Ningxia and Tianjin) were omitted because not enough closed forest could be found outside the reserve borders for us to evaluate effectiveness. For the unmatched method, we sampled n_1_ pixels of forest in year 2000 inside an NNR, and n_2_ pixels outside the NNRs (≥ 10 km and ≤ 100 km from the reserve borders). n_1_ = min(2000, 50%*N_1_), where N_1_ is the total number of forest pixels inside all NNRs within a province. n_2_ = min(10000, 50%*N_2_)), where N_2_ is the total number of forest pixels outside all of the NNRs within a province. The 10 km buffer was used to remove any leakage effects, in which deforestation banned in the reserve spills over to just outside the reserve (59, 60) or where deforestation is *lower* next to reserve borders, possibly due to governance or isolation spillovers (26, 61). Restricting pixels to within 100 km and to the same province was used to ensure that matching pixels were chosen from similar bioclimatic and governance regimes. Some of the n_1_ and n_2_ pixels were deforested (changed to neither forest nor woodland) in 2010. The effectiveness of NNRs within a province was calculated as Effectiveness.unmatched = deforestation.rate.outside – deforestation.rate.inside, where deforestation.rate.outside = (number of pixels converted to non-forest outside NNR)/n_2_ and deforestation.rate.nnr = (number of pixels converted to non-forest inside NNR)/n_1_. For the matched method (also known as ‘covariate matching’) (40), our four covariates were: elevation above sea level, TPI, slope, and distance to forest edge. For each within-NNR pixel, we found matching outside-NNR pixels (40, 62) within calipers (allowable differences) of ≤ 200 m of elevation, ≤ 15 m of TPI, and ≤ 5 degrees of slope. The outside-NNR pixel with the shortest Mahanolobis distance was deemed the best match (40). For the provincial-level analyses, we resampled 10 times and used the mean. For the individual NNR analyses, we resampled 20 times and used the mean. The effectiveness of NNRs within a province was calculated as Effectiveness.matched = (mean deforestation rate of matched samples outside all NNRs) - (mean deforestation rate of matched samples within all NNRs). Further methodological details are in SI Text.

### Author contributions

JGZ, GPR, SSY, and DWY designed the study. GPR, SSY, LW produced the land cover and forest change datasets. WW and JSL provided the national nature reserve dataset. SSY and DWY wrote the first draft, and all authors made extensive contributions to the text.

## Acknowledgments

This study was supported by Yunnan Province (20080A001), the Chinese Academy of Sciences (0902281081, KSCX2-YW-Z-1027), the National Natural Science Foundation of China (31272327, 31170498), the Ministry of Environmental Protection of China (20120928), the Ministry of Science and Technology of China (2012FY110800), the University of East Anglia, and the State Key Laboratory of Genetic Resources and Evolution at the Kunming Institute of Zoology, Salem State University (SSU). We thank Matthew Clark for guidance on *randomForests* and Tommy O’Connell, Sujan Joshi, and Zhenyang Hua for assistance with digitizing Google Earth ground truth data. Without the free dataset and software from NASA Land Processes Distributed Active Archive Center (LP DAAC), USGS/Earth Resources Observation and Science (EROS) Center, Sioux Falls, South Dakota (https://lpdaac.usgs.gov), there would be no study.

## Supplemental Material

### Supporting Text 1. Methods

#### SI 1.1 Terrain data and processing

The Shuttle Radar Topography Mission (SRTM) dataset was downloaded from NASA (earthexplorer.usgs.gov, accessed during May-November 2012) and reprojected to the Albers conic equal area projection for all 34 regions at 90 m resolution. Elevation, slope and topographic position index (TPI) were derived from the 90 m resolution SRTM using the raster package (1) in R v. 2.15 (2). The TPI of a pixel was defined as the difference between the elevation and the focal mean using a 11 by 11 Gaussian filter (*F*, eq. 1–2).

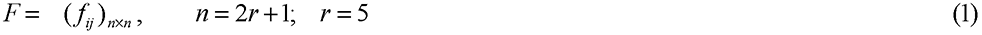

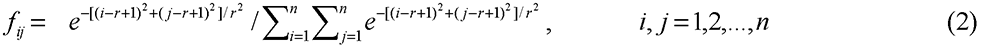

#### SI 1.2 Remote Sensing data and processing

We employed MOD13Q1 to map land cover and deforestation over mainland China from 2000-2010, because this global dataset allows consistent spatial and temporal comparison of vegetation conditions since February 2000 (3). For each year there are 23 scenes, from day 001 to day 353. Year 2000, however, did not start processing data until scene 049 (missing 001, 017, 033). Each scene of the MOD13Q1 dataset has 12 layers, including a 16-day composite of red (620–670 nm), near infrared (NIR: 841–876 nm), and mid-infrared (MIR: 2105–2155 nm) reflectance, Normalized Difference Vegetation Index (NDVI), and Enhanced Vegetation Index (EVI) at 231.7 m resolution (3). To cover all of mainland China for each scene, 19 tiles (H23V04-05, H24V04-06, H25V03-06, H26V03-06, H27V04-06, and H28V05-07) of MOD13Q1 data were needed (Figure S1). For all 19 tiles, we downloaded 274 scenes (from 2000-049, 2000-065, …, 2011-353, 2012-001) from the NASA Echo Reverb Data Portal (reverb.echo.nasa.gov/reverb/) in June 2012. Low-quality pixels of five layers (NDVI, EVI, MIR, NIR, red) in each scene of MOD13Q1 data, due to clouds and shadows, were filled by the mean values of the previous and the next scenes using the *raster* package (1) in *R*, thus achieving a filled MOD13Q1 dataset.

We use the 1:4,000,000 border of mainland China from the National Geomatics Center of China (ngcc.sbsm.gov.cn, accessed in May 22, 2003) and reprojected to an Albers conic equal area projection (WGS Albers 1984 projection, WGS84 datum with the standard parallels 25N, 47N, and central meridian 110E), which is the common projection used with geographic data across China. Therefore, mainland China was partitioned into 34 regions for land cover mapping and change detection (Figure S2). Then we reprojected and mosaiced all 274 scenes of filled MOD13Q1 data for each of the 34 regions to the same Albers conic equal area projection using nearest-neighbor resampling. All these processing were conducted using the *raster* package and the MODIS Reprojection Tool (lpdaac.usgs.gov/tools/modis_reprojection_tool, accessed in May 15, 2012).

**Figure S1.**
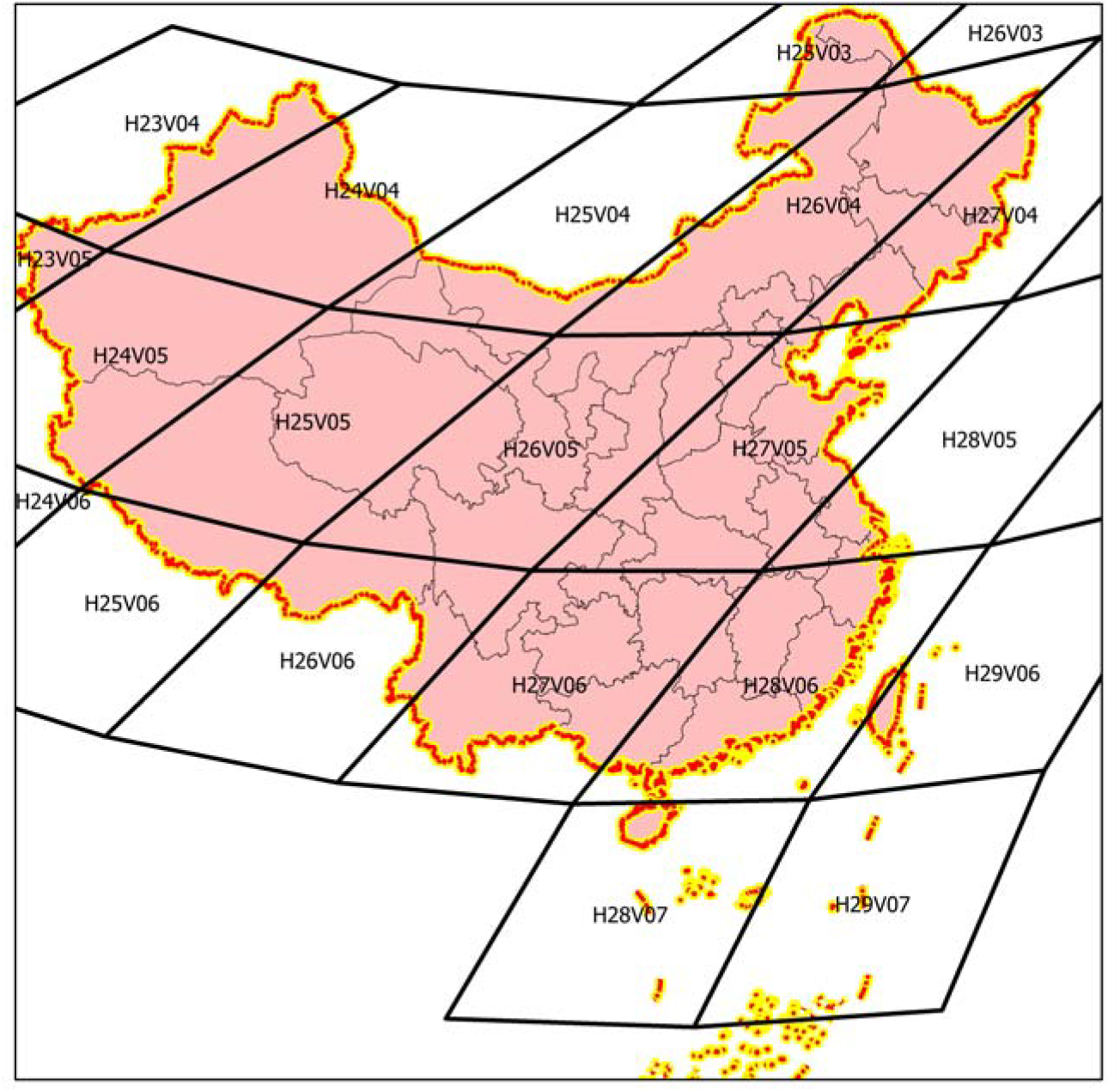
The 19 tiles of MOD13Q1 covering mainland China.

**Figure S2.**
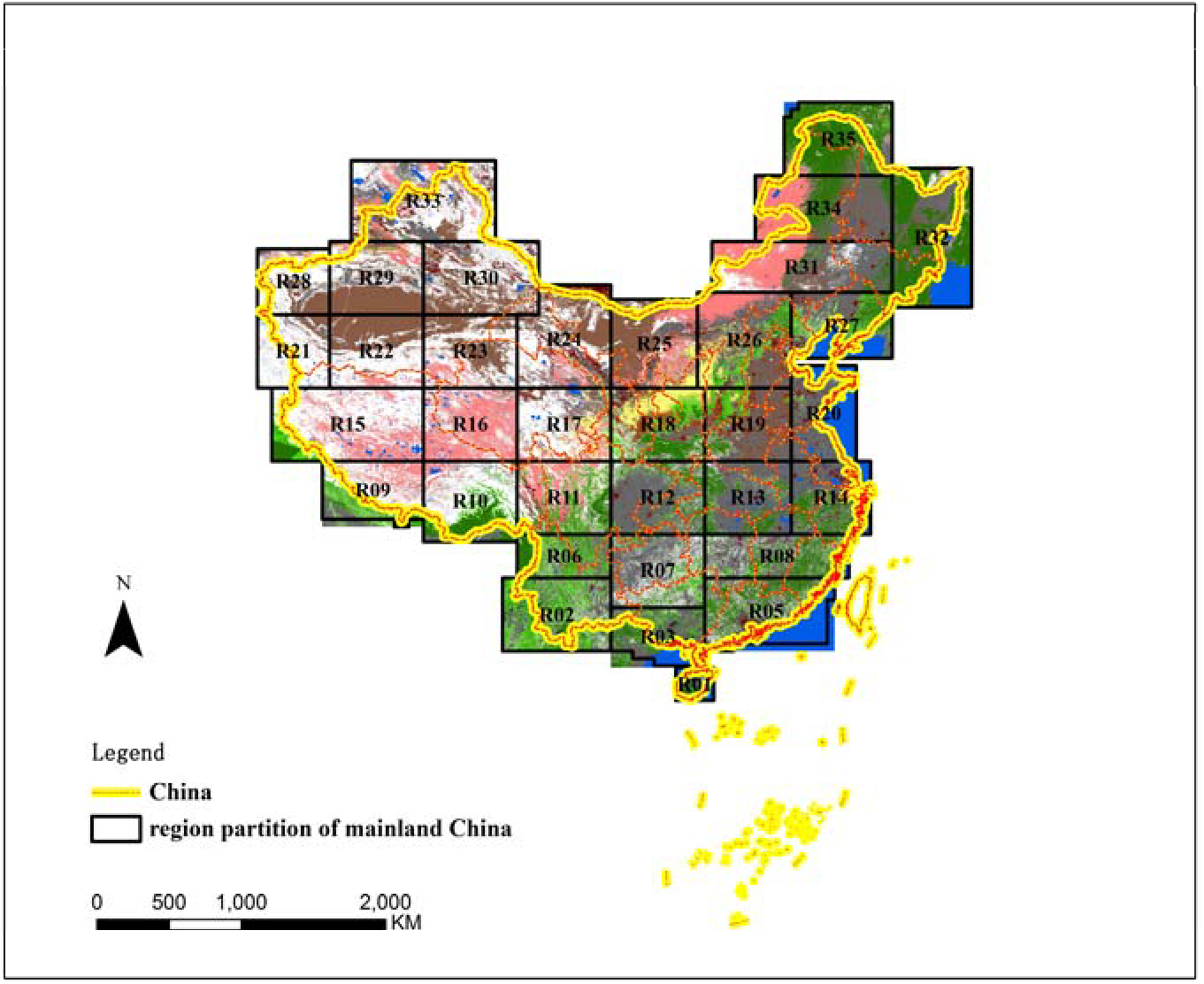
Partitioning of mainland China for land cover mapping.

#### SI 1.3 Land cover mapping and deforestation detection

##### Classification scheme

We define the training data for classification (4). We classified all of mainland China into (1) non-wooded land (tree canopy cover <40% and tree + shrub cover < 60%, or rubber plantation), and (2) forest (tree canopy cover ≥ 70%) and woodland (40%: S tree canopy cover < 70% and tree + shrub cover ≥ 60%). Forest and woodland included conifer forest and woodland, broadleaved forest and woodland, and mixed forest and woodland. Non-wooded land included urban (built-up land), agricultural land, grassland, open-water, perennial snow or ice, etc. Rubber plantations, with a regular canopy texture in high resolution images, could be distinguished from other forests and woodland. Therefore, we combined rubber plantation with non-wooded land in this study.

##### Training set definition

Training reference data came from high resolution imagery in Google Earth (GE, www.earth.google.com). GE now provides extensive coverage of China with high resolution imagery from DigitalGlobe. Many land cover studies adjudge GE to be accurate reference data (5–7). GE data were also used for some post-classification accuracy assessments. We accessed GE in ArcGIS software (V. 10.0, ESRI) through the ArcGIS plug-in Arc2Earth (www.arc2earth.com, last accessed in November 14, 2013).

By visual interpretation of high resolution images from GE, we digitized 13,869 polygons (with a minimal edge wider than 500 m) across mainland China and environs, labeling as forest (tree canopy cover ≥ 70%), woodland (40% ≤ tree canopy cover < 70% and tree + shrub cover ≥ 60%), or non-wooded land (tree canopy cover <40% and tree + shrub cover < 60%, or rubber plantation). These polygons were also labeled with detailed land cover types (such as conifer forest). We took these polygons as “ground-truth” data, as other studies have done (6, 7). Within each cell of one degree in longitude and one degree in latitude, we attempted to obtain 5 or more forest and woodland polygons if possible (Figure S3).

**Figure S3.**
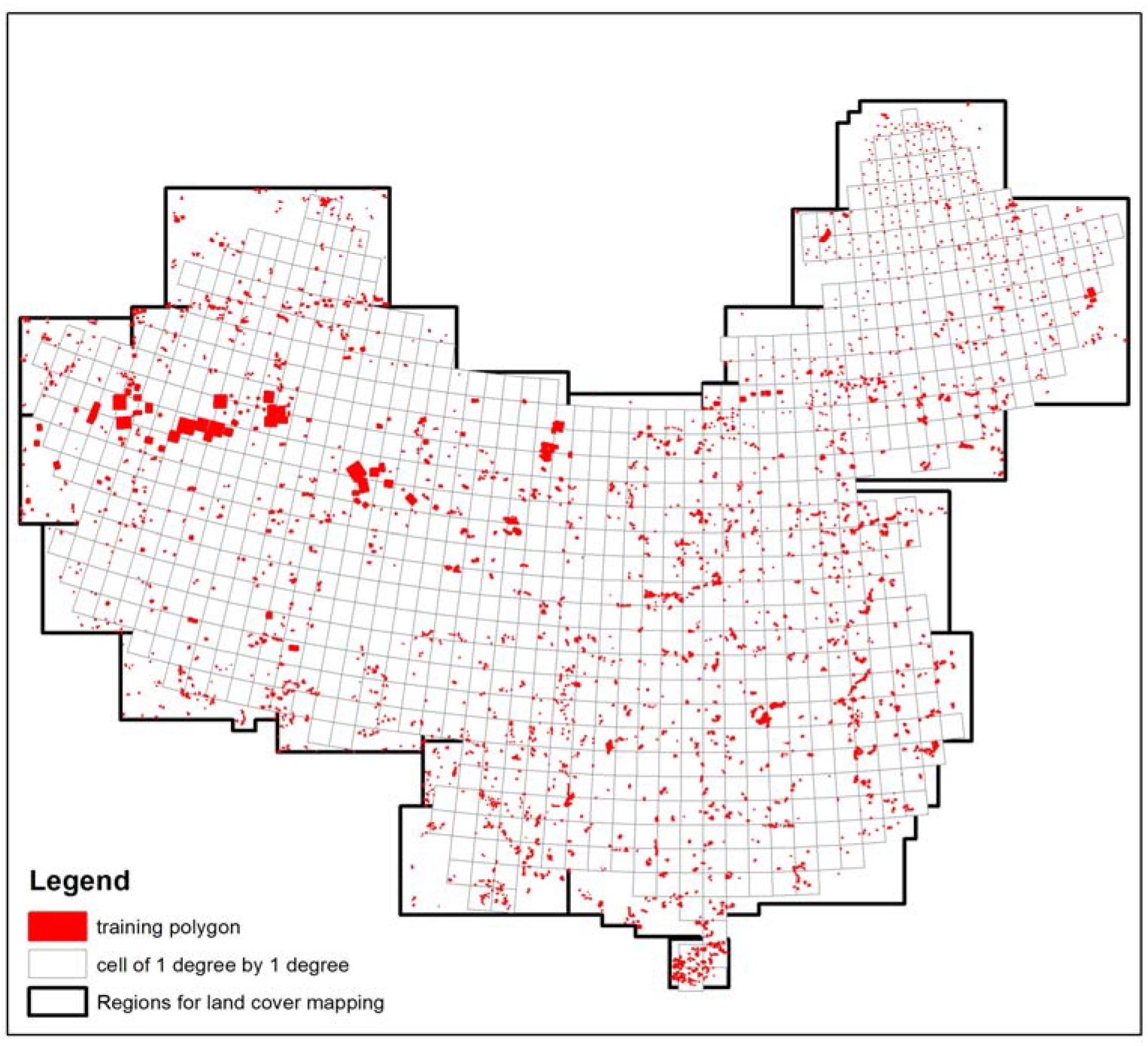
Training data by visual interpretation of high resolution images.

##### Land-cover mapping

We mapped land cover across mainland China region by region for each year from 2000-2011 using the *randomForest* algorithm (7, 8). For each year, 23 scenes of the five filled MOD13Q1 layers (NDVI, EVI, MIR, NIR, and red) were used as input except the year 2000 (only 19 filled scenes). The GE “ground-truth” data were used as training data for all years. These training data are imperfect because a few pixels might have changed from 2000-2010, and it is infeasible to verify the training data in the field. We performed land cover classification using the default parameters of the randomForest function in the *RandomForest* package (9). The producer’s accuracies of land cover maps in all 34 regions in all 12 years were higher than 92%.

##### Deforestation detection

We used three-year moving windows to adjudge forest loss (forest to non-wooded land). Only if the first year of a window was forest and the next two years were non-forest and non-woodland was a pixel classed as having been deforested in that window. To assess deforestation change-detection accuracy, we sampled 200 random deforested pixels in four regions (two in southern China, one in eastern China and one in Northeastern China), and these were checked with Google Earth. The user’s accuracy of deforestation detection was higher than 90% in all four regions, which demonstrated good change detection.

#### SI 1.4 Evaluation of the effectiveness of National-level Nature Reserves (NNRs)

##### NNR data

One mission of the Ministry of Environmental Protection (MEP) of the People’s Republic of China is to “Guide, coordinate and supervise the environmental protection of various kinds of nature reserves” (english.mep.gov.cn/About_SEPA/Mission/200803/t20080318_119444.htm), which includes archiving and monitoring the management plans of all NNRs. The Chinese Research Academy of Environmental Sciences (CRAES) was authorized by the MEP to use the management plans to digitize the boundaries of all NNRs. We evaluated the effectiveness of NNRs in protecting forest for each NNR and each province by comparing deforestation rates inside and outside NNRs, using both unmatched and matched approaches.

##### Unmatched approach

To estimate the deforestation rate of each of the 363 NNRs, we sampled *N_inside_* forested pixels inside each NNR in the year 2000 (Eq. 3). We also sampled *N_outside_* pixels outside each NNR (in a buffer zone ≥ 10 km from all NNRs and ≤ 100 km from the NNR) (Eq. 4). Let *n_inside_* and *n_outside_*be the *N_inside_* and *N_outside_* pixels, respectively, that changed to non-wooded land in the year 2010. The effectiveness of an NNR in preventing forest loss during 2000-2010 is given by Equation 5. We call this the “unmatched approach”.

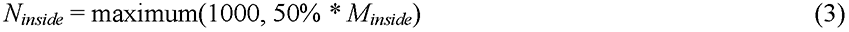

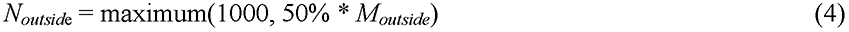

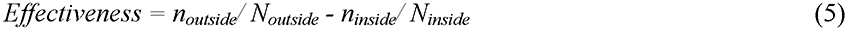

We only evaluated NNRs with more than 10 km^2^ forest cover in the year 2000 both inside and outside the NNR (both *N_inside_* and *N_outside_*>186 = 10/0.2317*0.2317). We therefore analyzed only 237 of 363 NNRs. For each analyzed NNR, we repeated this procedure 20 times.

At the province level, we sampled at most 2000 or 50% of the within-NNR-forested pixels in the year 2000 within each province. We also sampled at most 10,000 or 50% of the outside-NNR-forested pixels in the year 2000 within each province, in a buffer zone ≥ 10 km and ≤ 100 km from all NNR borders within each province. 26 of 31 provinces could be so analyzed, omitting Shanghai, Tianjin, Ningxia, Jiangsu, and Shandong because of too few forest pixels outside the NNRs. For each of the remaining 26 provinces, we repeated this procedure 10 times.

##### Matched approach

The unmatched approach assumes that NNR borders have been sited randomly, which is untrue (10). To control for environmental differences between pixels inside NNRs and outside NNRs, we also employed a matched approach, also known as covariate matching (10, 11). Our covariates included elevation above sea level, TPI (topographic position index), slope, and distance to forest edge. For each sample pixel from an NNR, 1) we found all matching pixels outside the NNR within calipers (allowable differences) of ≤ 200 m of elevation, ≤ 15 m of TPI, and ≤ 5 degrees of slope; and 2) of these pixels, we chose the inside pixel’s match as the outside pixel with the shortest Mahanolobis distance (10). We repeated this procedure 20 times per NNR. We also repeated the analysis at the provincial level, pooling all NNRs within a province and repeating the procedure 10 times.

The matched approach is considered a more reliable gauge of NNR protective efficacy (10, 12) because it partials out the confounding effect of protection by terrain inaccessibility *per se*, but since it is not yet possible in China to avoid sampling pixels in provincial-municipal- and county-level nature reserves, which also receive some degree of protection, the matched approach is likely to be somewhat conservative. We thus consider both methods in order to bracket the true effectiveness of NNR protection.

### Supporting Text 2. Results

**Table S1.** Effectiveness of the 237 NNRs in preventing deforestation. **Prov**: province key for each of the 237 NNRs. See Main Text Figure 1 legend for translation to full province names. **Defor. NNR**: annual deforestation rate inside the NNR. **Eff. Unmat**.: effectiveness of the NNR evaluated by unmatched approach. **Eff. mat**.: effectiveness of the NNR evaluated by the matched approach. **Sample size inside/outside**: the number of sampled pixels inside and outside the NNR. **Sample matched rate**: percentage of sampled pixels inside the NNR which could be matched to a pixel outside the NNR. **p-value unm**.: the possibility of the effectiveness of the NNR evaluated by unmatched approach that is equal to zero by t-test (N = 20). **p-value unm**.: the possibility of the effectiveness of the NNR evaluated by matched approach that is equal to zero by t-test (N = 20).

| Prov. | Deform. NNR | Eff. Unmat. | Eff. Mat. | Sample size inside | Sample size outside | Sample matched rate | p-value unnm. | p-value mat. |
| --- | --- | --- | --- | --- | --- | --- | --- | --- |
| AH | 0.061 | 0.048 | 0.023 | 1000 | 10000 | 0.966 | 0.000 | 0.000 |
| AH | 0.057 | 0.053 | 0.020 | 1000 | 10000 | 0.918 | 0.000 | 0.000 |
| AH | 0.041 | 0.068 | 0.018 | 589 | 10000 | 0.899 | 0.000 | 0.000 |
| AH | 0.197 | -0.086 | -0.062 | 471 | 10000 | 0.883 | 0.000 | 0.000 |
| BJ | 0.007 | 0.024 | -0.004 | 1000 | 10000 | 0.840 | 0.000 | 0.000 |
| BJ | 0.000 | 0.031 | 0.001 | 375 | 10000 | 0.920 | 0.000 | 0.008 |
| CQ | 0.034 | 0.103 | 0.036 | 1000 | 10000 | 0.961 | 0.000 | 0.000 |
| CQ | 0.027 | 0.110 | 0.017 | 1000 | 10000 | 0.988 | 0.000 | 0.000 |
| CQ | 0.116 | 0.020 | -0.043 | 1000 | 10000 | 0.886 | 0.000 | 0.000 |
| CQ | 0.167 | -0.032 | -0.035 | 801 | 10000 | 0.924 | 0.000 | 0.000 |
| CQ | 0.031 | 0.105 | 0.100 | 323 | 10000 | 1.000 | 0.000 | 0.000 |
| FJ | 0.002 | 0.145 | 0.050 | 1000 | 10000 | 0.970 | 0.000 | 0.000 |
| FJ | 0.005 | 0.140 | 0.022 | 1000 | 10000 | 0.856 | 0.000 | 0.000 |
| FJ | 0.264 | -0.117 | -0.096 | 1000 | 10000 | 1.000 | 0.000 | 0.000 |
| FJ | 0.003 | 0.144 | 0.038 | 1000 | 9999 | 0.997 | 0.000 | 0.000 |
| FJ | 0.023 | 0.123 | 0.085 | 1000 | 9999 | 0.999 | 0.000 | 0.000 |
| FJ | 0.101 | 0.046 | -0.005 | 1000 | 10000 | 1.000 | 0.000 | 0.097 |
| FJ | 0.030 | 0.116 | -0.003 | 1000 | 10000 | 0.961 | 0.000 | 0.192 |
| FJ | 0.032 | 0.115 | 0.051 | 1000 | 10000 | 0.974 | 0.000 | 0.000 |
| FJ | 0.008 | 0.140 | 0.117 | 972 | 10000 | 1.000 | 0.000 | 0.000 |
| FJ | 0.005 | 0.143 | 0.028 | 833 | 10000 | 0.994 | 0.000 | 0.000 |
| GD | 0.036 | 0.363 | 0.069 | 1000 | 9998 | 0.923 | 0.000 | 0.000 |
| GD | 0.087 | 0.311 | 0.114 | 1000 | 9998 | 0.964 | 0.000 | 0.000 |
| GD | 0.211 | 0.187 | 0.002 | 846 | 9998 | 0.999 | 0.000 | 0.633 |
| GD | 0.028 | 0.369 | 0.190 | 649 | 9999 | 0.998 | 0.000 | 0.000 |
| GD | 0.417 | -0.021 | 0.037 | 211 | 9999 | 1.000 | 0.001 | 0.000 |
| GS | 0.183 | -0.064 | -0.045 | 1000 | 10000 | 0.982 | 0.000 | 0.000 |
| GS | 0.222 | -0.103 | 0.259 | 1000 | 10000 | 0.928 | 0.000 | 0.000 |
| GS | 0.410 | -0.291 | 0.088 | 1000 | 10000 | 0.930 | 0.000 | 0.000 |
| GS | 0.277 | -0.156 | 0.221 | 1000 | 10000 | 0.925 | 0.000 | 0.000 |
| GS | 0.033 | 0.087 | 0.020 | 1000 | 10000 | 0.941 | 0.000 | 0.000 |
| GS | 0.276 | -0.155 | 0.196 | 887 | 10000 | 0.978 | 0.000 | 0.000 |
| GS | 0.545 | -0.426 | -0.084 | 506 | 10000 | 0.998 | 0.000 | 0.000 |
| GS | 0.625 | -0.506 | -0.002 | 264 | 10000 | 0.913 | 0.000 | 0.841 |
| GX | 0.207 | 0.152 | -0.012 | 1000 | 10000 | 0.904 | 0.000 | 0.073 |
| GX | 0.064 | 0.298 | 0.147 | 1000 | 9999 | 0.989 | 0.000 | 0.000 |
| GX | 0.007 | 0.354 | 0.176 | 1000 | 10000 | 0.926 | 0.000 | 0.000 |
| GX | 0.040 | 0.320 | 0.120 | 1000 | 9999 | 0.991 | 0.000 | 0.000 |
| GX | 0.040 | 0.320 | 0.183 | 1000 | 10000 | 0.990 | 0.000 | 0.000 |
| GX | 0.019 | 0.342 | 0.247 | 1000 | 10000 | 0.995 | 0.000 | 0.000 |
| GX | 0.002 | 0.360 | 0.078 | 973 | 10000 | 0.888 | 0.000 | 0.000 |
| GX | 0.000 | 0.362 | 0.298 | 919 | 9999 | 0.998 | 0.000 | 0.000 |
| GX | 0.084 | 0.275 | 0.082 | 909 | 9999 | 0.947 | 0.000 | 0.000 |
| GX | 0.006 | 0.355 | 0.072 | 865 | 10000 | 0.927 | 0.000 | 0.000 |
| GX | 0.040 | 0.320 | 0.267 | 623 | 10000 | 0.996 | 0.000 | 0.000 |
| GX | 0.531 | -0.168 | -0.137 | 443 | 10000 | 0.954 | 0.000 | 0.000 |
| GX | 0.578 | -0.217 | -0.312 | 312 | 10000 | 0.975 | 0.000 | 0.000 |
| GX | 0.296 | 0.065 | 0.123 | 141 | 9999 | 1.000 | 0.000 | 0.000 |
| GZ | 0.047 | 0.251 | 0.036 | 1000 | 10000 | 0.938 | 0.000 | 0.000 |
| GZ | 0.013 | 0.287 | 0.178 | 1000 | 10000 | 0.987 | 0.000 | 0.000 |
| GZ | 0.353 | -0.054 | -0.189 | 1000 | 10000 | 0.992 | 0.000 | 0.000 |
| GZ | 0.092 | 0.207 | 0.120 | 1000 | 10000 | 0.997 | 0.000 | 0.000 |
| GZ | 0.003 | 0.295 | 0.023 | 1000 | 10000 | 0.974 | 0.000 | 0.000 |
| GZ | 0.009 | 0.290 | 0.120 | 825 | 10000 | 0.998 | 0.000 | 0.000 |
| GZ | 0.100 | 0.198 | 0.159 | 466 | 10000 | 0.953 | 0.000 | 0.000 |
| HA | 0.031 | 0.085 | 0.023 | 1000 | 10000 | 0.998 | 0.000 | 0.000 |
| HA | 0.132 | -0.015 | 0.000 | 1000 | 10000 | 0.999 | 0.000 | 0.895 |
| HA | 0.014 | 0.102 | 0.014 | 1000 | 10000 | 0.972 | 0.000 | 0.000 |
| HA | 0.079 | 0.038 | 0.008 | 1000 | 10000 | 1.000 | 0.000 | 0.013 |
| HA | 0.108 | 0.009 | 0.026 | 1000 | 10000 | 0.999 | 0.000 | 0.000 |
| HA | 0.034 | 0.081 | 0.038 | 1000 | 10000 | 0.964 | 0.000 | 0.000 |
| HA | 0.107 | 0.011 | 0.043 | 558 | 10000 | 0.997 | 0.000 | 0.000 |
| HA | 0.162 | -0.045 | -0.057 | 379 | 10000 | 1.000 | 0.000 | 0.000 |
| HA | 0.073 | 0.043 | -0.002 | 347 | 10000 | 0.933 | 0.000 | 0.739 |
| HB | 0.037 | 0.092 | 0.022 | 1000 | 10000 | 1.000 | 0.000 | 0.000 |
| HB | 0.118 | 0.012 | -0.025 | 1000 | 10000 | 0.977 | 0.000 | 0.000 |
| HB | 0.056 | 0.073 | 0.047 | 1000 | 10000 | 0.979 | 0.000 | 0.000 |
| HB | 0.134 | -0.005 | -0.006 | 1000 | 10000 | 1.000 | 0.103 | 0.065 |
| HB | 0.038 | 0.091 | 0.013 | 1000 | 10000 | 1.000 | 0.000 | 0.000 |
| HB | 0.027 | 0.102 | 0.024 | 1000 | 10000 | 0.753 | 0.000 | 0.000 |
| HB | 0.055 | 0.074 | 0.013 | 1000 | 10000 | 0.988 | 0.000 | 0.000 |
| HB | 0.163 | -0.033 | -0.021 | 1000 | 10000 | 0.998 | 0.000 | 0.000 |
| HE | 0.005 | 0.031 | 0.006 | 1000 | 10000 | 0.998 | 0.000 | 0.000 |
| HE | 0.009 | 0.026 | 0.008 | 1000 | 10000 | 1.000 | 0.000 | 0.000 |
| HE | 0.006 | 0.030 | 0.000 | 1000 | 10000 | 1.000 | 0.000 | 1.000 |
| HE | 0.054 | -0.018 | 0.009 | 1000 | 10000 | 0.705 | 0.000 | 0.003 |
| HE | 0.000 | 0.036 | 0.004 | 815 | 10000 | 0.999 | 0.000 | 0.000 |
| HE | 0.091 | -0.056 | 0.017 | 602 | 10000 | 0.999 | 0.000 | 0.054 |
| HE | 0.000 | 0.036 | 0.005 | 429 | 10000 | 0.993 | 0.000 | 0.000 |
| HE | 0.129 | -0.093 | 0.001 | 376 | 10000 | 1.000 | 0.000 | 0.864 |
| HE | 0.000 | 0.036 | 0.011 | 144 | 10000 | 0.967 | 0.000 | 0.000 |
| HE | 0.084 | -0.048 | -0.052 | 93 | 10000 | 0.996 | 0.000 | 0.000 |
| HI | 0.026 | 0.148 | 0.017 | 1000 | 7820 | 0.998 | 0.000 | 0.000 |
| HI | 0.004 | 0.171 | 0.017 | 1000 | 7820 | 0.999 | 0.000 | 0.000 |
| HI | 0.004 | 0.171 | 0.026 | 1000 | 7818 | 0.997 | 0.000 | 0.000 |
| HI | 0.006 | 0.170 | 0.002 | 1000 | 7821 | 0.951 | 0.000 | 0.047 |
| HL | 0.012 | 0.049 | 0.040 | 1000 | 10000 | 1.000 | 0.000 | 0.000 |
| HL | 0.062 | -0.001 | -0.011 | 1000 | 10000 | 1.000 | 0.730 | 0.013 |
| HL | 0.231 | -0.169 | -0.110 | 1000 | 10000 | 1.000 | 0.000 | 0.000 |
| HL | 0.003 | 0.058 | 0.023 | 1000 | 10000 | 1.000 | 0.000 | 0.000 |
| HL | 0.032 | 0.029 | -0.004 | 1000 | 10000 | 1.000 | 0.000 | 0.017 |
| HL | 0.014 | 0.047 | 0.051 | 1000 | 10000 | 1.000 | 0.000 | 0.000 |
| HL | 0.007 | 0.055 | 0.014 | 1000 | 10000 | 1.000 | 0.000 | 0.000 |
| HL | 0.022 | 0.039 | -0.005 | 1000 | 10000 | 0.999 | 0.000 | 0.010 |
| HL | 0.001 | 0.060 | 0.017 | 1000 | 10000 | 1.000 | 0.000 | 0.000 |
| HL | 0.008 | 0.053 | 0.002 | 1000 | 10000 | 1.000 | 0.000 | 0.028 |
| HL | 0.001 | 0.060 | 0.004 | 1000 | 10000 | 1.000 | 0.000 | 0.000 |
| HL | 0.085 | -0.024 | 0.014 | 1000 | 10000 | 1.000 | 0.000 | 0.008 |
| HL | 0.069 | -0.008 | -0.013 | 1000 | 10000 | 1.000 | 0.002 | 0.000 |
| HL | 0.035 | 0.026 | -0.001 | 1000 | 10000 | 1.000 | 0.000 | 0.824 |
| HL | 0.029 | 0.032 | 0.022 | 1000 | 10000 | 1.000 | 0.000 | 0.000 |
| HL | 0.128 | -0.067 | 0.097 | 1000 | 10000 | 1.000 | 0.000 | 0.000 |
| HL | 0.009 | 0.052 | 0.004 | 1000 | 10000 | 1.000 | 0.000 | 0.000 |
| HL | 0.059 | 0.002 | 0.013 | 1000 | 10000 | 1.000 | 0.186 | 0.000 |
| HL | 0.035 | 0.026 | 0.028 | 1000 | 10000 | 1.000 | 0.000 | 0.000 |
| HL | 0.334 | -0.273 | -0.163 | 540 | 10000 | 1.000 | 0.000 | 0.000 |
| HL | 0.404 | -0.343 | -0.047 | 170 | 10000 | 1.000 | 0.000 | 0.024 |
| HL | 0.146 | -0.085 | -0.080 | 69 | 10000 | 1.000 | 0.000 | 0.000 |
| HN | 0.043 | 0.288 | 0.244 | 1000 | 10000 | 0.941 | 0.000 | 0.000 |
| HN | 0.163 | 0.170 | 0.118 | 1000 | 10000 | 0.998 | 0.000 | 0.000 |
| HN | 0.044 | 0.288 | 0.211 | 1000 | 10000 | 0.895 | 0.000 | 0.000 |
| HN | 0.057 | 0.276 | 0.262 | 1000 | 10000 | 0.995 | 0.000 | 0.000 |
| HN | 0.019 | 0.311 | 0.107 | 1000 | 10000 | 0.990 | 0.000 | 0.000 |
| HN | 0.041 | 0.290 | 0.067 | 1000 | 10000 | 1.000 | 0.000 | 0.000 |
| HN | 0.085 | 0.247 | 0.183 | 1000 | 10000 | 0.996 | 0.000 | 0.000 |
| HN | 0.010 | 0.322 | 0.118 | 1000 | 10000 | 0.913 | 0.000 | 0.000 |
| HN | 0.030 | 0.300 | 0.122 | 1000 | 10000 | 0.994 | 0.000 | 0.000 |
| HN | 0.086 | 0.246 | 0.292 | 1000 | 10000 | 1.000 | 0.000 | 0.000 |
| HN | 0.113 | 0.217 | 0.033 | 1000 | 10000 | 0.908 | 0.000 | 0.000 |
| HN | 0.063 | 0.269 | 0.286 | 1000 | 10000 | 0.993 | 0.000 | 0.000 |
| HN | 0.221 | 0.109 | -0.013 | 946 | 10000 | 0.993 | 0.000 | 0.003 |
| HN | 0.157 | 0.172 | 0.047 | 828 | 10000 | 0.969 | 0.000 | 0.000 |
| HN | 0.007 | 0.324 | 0.112 | 723 | 10000 | 0.906 | 0.000 | 0.000 |
| HN | 0.404 | -0.071 | 0.039 | 477 | 10000 | 1.000 | 0.000 | 0.000 |
| HN | 0.448 | -0.119 | -0.022 | 224 | 10000 | 1.000 | 0.000 | 0.030 |
| JL | 0.008 | 0.025 | 0.009 | 1000 | 10000 | 1.000 | 0.000 | 0.000 |
| JL | 0.030 | 0.002 | -0.009 | 1000 | 10000 | 1.000 | 0.039 | 0.000 |
| JL | 0.042 | -0.009 | 0.036 | 1000 | 10000 | 1.000 | 0.000 | 0.000 |
| JL | 0.007 | 0.025 | 0.003 | 1000 | 10000 | 0.893 | 0.000 | 0.016 |
| JL | 0.013 | 0.020 | 0.007 | 1000 | 10000 | 1.000 | 0.000 | 0.001 |
| JL | 0.013 | 0.020 | 0.009 | 1000 | 10000 | 1.000 | 0.000 | 0.001 |
| JL | 0.027 | 0.005 | 0.009 | 1000 | 10000 | 1.000 | 0.000 | 0.000 |
| JL | 0.007 | 0.026 | 0.002 | 1000 | 10000 | 0.999 | 0.000 | 0.081 |
| JL | 0.057 | -0.024 | -0.041 | 1000 | 10000 | 0.996 | 0.000 | 0.000 |
| JL | 0.057 | -0.024 | 0.041 | 1000 | 10000 | 1.000 | 0.000 | 0.000 |
| JX | 0.032 | 0.132 | 0.000 | 1000 | 10000 | 0.997 | 0.000 | 0.888 |
| JX | 0.093 | 0.070 | 0.045 | 1000 | 10000 | 0.997 | 0.000 | 0.000 |
| JX | 0.005 | 0.159 | 0.030 | 1000 | 10000 | 0.650 | 0.000 | 0.000 |
| JX | 0.013 | 0.150 | 0.032 | 1000 | 10000 | 0.942 | 0.000 | 0.000 |
| JX | 0.022 | 0.143 | 0.059 | 1000 | 10000 | 0.999 | 0.000 | 0.000 |
| JX | 0.004 | 0.160 | 0.026 | 1000 | 10000 | 0.965 | 0.000 | 0.000 |
| JX | 0.101 | 0.064 | 0.080 | 1000 | 10000 | 1.000 | 0.000 | 0.000 |
| JX | 0.006 | 0.159 | 0.030 | 1000 | 10000 | 0.976 | 0.000 | 0.000 |
| JX | 0.011 | 0.154 | 0.048 | 995 | 10000 | 0.998 | 0.000 | 0.000 |
| LN | 0.001 | 0.052 | 0.017 | 1000 | 10000 | 0.999 | 0.000 | 0.000 |
| LN | 0.037 | 0.016 | 0.117 | 955 | 10000 | 1.000 | 0.000 | 0.000 |
| LN | 0.000 | 0.053 | 0.016 | 695 | 10000 | 0.999 | 0.000 | 0.000 |
| LN | 0.243 | -0.190 | -0.102 | 221 | 10000 | 1.000 | 0.000 | 0.000 |
| LN | 0.140 | -0.087 | -0.047 | 114 | 10000 | 0.996 | 0.000 | 0.000 |
| NM | 0.065 | 0.007 | 0.029 | 1000 | 10000 | 1.000 | 0.001 | 0.000 |
| NM | 0.004 | 0.068 | 0.024 | 1000 | 10000 | 1.000 | 0.000 | 0.000 |
| NM | 0.088 | -0.017 | 0.024 | 1000 | 10000 | 1.000 | 0.000 | 0.000 |
| NM | 0.176 | -0.105 | -0.077 | 1000 | 10000 | 0.996 | 0.000 | 0.000 |
| NM | 0.005 | 0.067 | 0.050 | 1000 | 10000 | 0.999 | 0.000 | 0.000 |
| NM | 0.024 | 0.049 | 0.063 | 1000 | 10000 | 1.000 | 0.000 | 0.000 |
| NM | 0.115 | -0.043 | -0.074 | 1000 | 10000 | 0.007 | 0.000 | 0.526 |
| NM | 0.184 | -0.112 | 0.052 | 603 | 10000 | 0.588 | 0.000 | 0.001 |
| NM | 0.185 | -0.113 | 0.040 | 130 | 10000 | 0.998 | 0.000 | 0.007 |
| QH | 0.430 | -0.104 | 0.181 | 1000 | 10000 | 0.933 | 0.000 | 0.000 |
| QH | 0.469 | -0.141 | -0.178 | 111 | 10000 | 0.840 | 0.000 | 0.000 |
| SC | 0.069 | 0.007 | 0.019 | 1000 | 10000 | 0.992 | 0.000 | 0.000 |
| SC | 0.000 | 0.076 | 0.008 | 1000 | 10000 | 0.996 | 0.000 | 0.000 |
| SC | 0.021 | 0.055 | 0.004 | 1000 | 10000 | 0.980 | 0.000 | 0.022 |
| SC | 0.015 | 0.061 | 0.009 | 1000 | 10000 | 0.990 | 0.000 | 0.000 |
| SC | 0.006 | 0.070 | 0.012 | 1000 | 10000 | 0.993 | 0.000 | 0.000 |
| SC | 0.050 | 0.026 | 0.126 | 1000 | 10000 | 0.947 | 0.000 | 0.000 |
| SC | 0.001 | 0.075 | 0.059 | 1000 | 10000 | 0.954 | 0.000 | 0.000 |
| SC | 0.295 | -0.219 | -0.219 | 1000 | 10000 | 0.991 | 0.000 | 0.000 |
| SC | 0.049 | 0.028 | 0.026 | 1000 | 10000 | 0.980 | 0.000 | 0.000 |
| SC | 0.064 | 0.013 | -0.027 | 1000 | 10000 | 0.986 | 0.000 | 0.000 |
| SC | 0.044 | 0.031 | 0.076 | 1000 | 10000 | 0.979 | 0.000 | 0.000 |
| SC | 0.133 | -0.056 | -0.021 | 1000 | 10000 | 0.990 | 0.000 | 0.000 |
| SC | 0.090 | -0.014 | -0.038 | 1000 | 10000 | 0.974 | 0.000 | 0.000 |
| SC | 0.438 | -0.362 | -0.294 | 1000 | 10000 | 0.973 | 0.000 | 0.000 |
| SC | 0.014 | 0.061 | 0.002 | 1000 | 10000 | 0.970 | 0.000 | 0.172 |
| SC | 0.127 | -0.050 | 0.066 | 752 | 10000 | 0.988 | 0.000 | 0.000 |
| SC | 0.012 | 0.065 | 0.041 | 615 | 10000 | 0.985 | 0.000 | 0.000 |
| SC | 0.744 | -0.668 | -0.610 | 580 | 10000 | 0.970 | 0.000 | 0.000 |
| SC | 0.176 | -0.099 | -0.043 | 531 | 10000 | 0.991 | 0.000 | 0.000 |
| SC | 0.022 | 0.054 | -0.007 | 312 | 10000 | 1.000 | 0.000 | 0.002 |
| SC | 0.007 | 0.068 | 0.039 | 295 | 10000 | 0.995 | 0.000 | 0.000 |
| SC | 0.021 | 0.054 | 0.010 | 273 | 10000 | 0.994 | 0.000 | 0.002 |
| SD | 0.398 | -0.317 | -0.179 | 694 | 2192 | 0.999 | 0.000 | 0.000 |
| SD | 0.046 | 0.033 | -0.007 | 467 | 2192 | 1.000 | 0.000 | 0.005 |
| SN | 0.012 | 0.019 | -0.001 | 1000 | 10000 | 0.992 | 0.000 | 0.470 |
| SN | 0.000 | 0.031 | 0.019 | 1000 | 10000 | 1.000 | 0.000 | 0.000 |
| SN | 0.036 | -0.005 | 0.005 | 1000 | 10000 | 1.000 | 0.000 | 0.018 |
| SN | 0.028 | 0.003 | 0.032 | 1000 | 10000 | 0.964 | 0.030 | 0.000 |
| SN | 0.141 | -0.110 | -0.100 | 1000 | 10000 | 0.958 | 0.000 | 0.000 |
| SN | 0.003 | 0.029 | 0.002 | 1000 | 10000 | 0.993 | 0.000 | 0.001 |
| SN | 0.050 | -0.019 | -0.012 | 1000 | 10000 | 0.980 | 0.000 | 0.000 |
| SN | 0.000 | 0.031 | 0.004 | 1000 | 10000 | 1.000 | 0.000 | 0.000 |
| SN | 0.064 | -0.033 | -0.008 | 1000 | 10000 | 0.732 | 0.000 | 0.011 |
| SN | 0.054 | -0.023 | -0.035 | 1000 | 10000 | 0.992 | 0.000 | 0.000 |
| SN | 0.000 | 0.031 | 0.006 | 1000 | 10000 | 1.000 | 0.000 | 0.000 |
| SN | 0.023 | 0.008 | -0.005 | 1000 | 10000 | 0.982 | 0.000 | 0.001 |
| SN | 0.011 | 0.019 | 0.000 | 1000 | 10000 | 0.989 | 0.000 | 0.627 |
| SN | 0.052 | -0.021 | -0.025 | 1000 | 10000 | 0.975 | 0.000 | 0.000 |
| SN | 0.055 | -0.024 | -0.038 | 580 | 10000 | 0.998 | 0.000 | 0.000 |
| SX | 0.007 | 0.006 | 0.006 | 1000 | 10000 | 0.996 | 0.000 | 0.000 |
| SX | 0.029 | -0.016 | -0.008 | 1000 | 10000 | 0.996 | 0.000 | 0.000 |
| SX | 0.005 | 0.007 | 0.001 | 982 | 10000 | 0.997 | 0.000 | 0.294 |
| SX | 0.000 | 0.013 | 0.001 | 940 | 10000 | 1.000 | 0.000 | 0.004 |
| SX | 0.000 | 0.013 | 0.006 | 518 | 10000 | 1.000 | 0.000 | 0.000 |
| SX | 0.012 | 0.000 | 0.051 | 468 | 10000 | 0.977 | 0.842 | 0.000 |
| XJ | 0.004 | 0.079 | 0.029 | 1000 | 10000 | 0.968 | 0.000 | 0.000 |
| XJ | 0.117 | -0.035 | -0.035 | 1000 | 10000 | 1.000 | 0.000 | 0.000 |
| XZ | 0.038 | -0.019 | -0.020 | 1000 | 10000 | 0.951 | 0.000 | 0.000 |
| XZ | 0.004 | 0.015 | 0.027 | 1000 | 10000 | 0.984 | 0.000 | 0.000 |
| XZ | 0.013 | 0.006 | -0.002 | 1000 | 10000 | 0.980 | 0.000 | 0.199 |
| XZ | 0.005 | 0.014 | 0.013 | 1000 | 10000 | 0.976 | 0.000 | 0.000 |
| XZ | 0.018 | 0.001 | 0.026 | 291 | 10000 | 0.965 | 0.548 | 0.000 |
| YN | 0.029 | 0.015 | -0.010 | 1000 | 10000 | 0.982 | 0.000 | 0.000 |
| YN | 0.011 | 0.031 | -0.003 | 1000 | 10000 | 0.978 | 0.000 | 0.005 |
| YN | 0.033 | 0.011 | -0.020 | 1000 | 10000 | 0.963 | 0.000 | 0.000 |
| YN | 0.130 | -0.087 | 0.004 | 1000 | 10000 | 0.965 | 0.000 | 0.260 |
| YN | 0.014 | 0.028 | 0.015 | 1000 | 10000 | 0.998 | 0.000 | 0.000 |
| YN | 0.030 | 0.013 | 0.013 | 1000 | 10000 | 0.998 | 0.000 | 0.000 |
| YN | 0.000 | 0.043 | 0.006 | 1000 | 10000 | 0.979 | 0.000 | 0.000 |
| YN | 0.032 | 0.012 | 0.093 | 1000 | 10000 | 0.996 | 0.000 | 0.000 |
| YN | 0.002 | 0.041 | 0.003 | 1000 | 10000 | 0.998 | 0.000 | 0.002 |
| YN | 0.138 | -0.094 | 0.048 | 1000 | 10000 | 0.910 | 0.000 | 0.000 |
| YN | 0.000 | 0.043 | 0.010 | 768 | 10000 | 1.000 | 0.000 | 0.000 |
| YN | 0.009 | 0.034 | 0.003 | 763 | 10000 | 1.000 | 0.000 | 0.014 |
| YN | 0.034 | 0.009 | 0.008 | 492 | 10000 | 0.885 | 0.000 | 0.003 |
| YN | 0.065 | -0.021 | -0.029 | 438 | 10000 | 1.000 | 0.000 | 0.000 |
| YN | 0.004 | 0.039 | 0.112 | 427 | 10000 | 0.991 | 0.000 | 0.000 |
| YN | 0.013 | 0.030 | 0.008 | 263 | 10000 | 0.936 | 0.000 | 0.002 |
| YN | 0.009 | 0.035 | 0.124 | 163 | 10000 | 0.996 | 0.000 | 0.000 |
| ZJ | 0.220 | -0.075 | -0.092 | 1000 | 9999 | 1.000 | 0.000 | 0.000 |
| ZJ | 0.073 | 0.072 | 0.078 | 1000 | 9999 | 0.997 | 0.000 | 0.000 |
| ZJ | 0.025 | 0.119 | 0.045 | 1000 | 9999 | 0.998 | 0.000 | 0.000 |
| ZJ | 0.081 | 0.064 | 0.058 | 952 | 9999 | 1.000 | 0.000 | 0.000 |
| ZJ | 0.006 | 0.138 | 0.053 | 917 | 9999 | 0.776 | 0.000 | 0.000 |
| ZJ | 0.002 | 0.143 | 0.018 | 517 | 9999 | 0.900 | 0.000 | 0.000 |
| ZJ | 0.102 | 0.042 | -0.004 | 289 | 9999 | 0.988 | 0.000 | 0.376 |
| ZJ | 0.232 | -0.088 | -0.135 | 191 | 9999 | 1.000 | 0.000 | 0.000 |

